# Z-Flipons conserved between human and mouse are associated with increased transcription initiation rates

**DOI:** 10.1101/2023.10.31.564984

**Authors:** Nazar Beknazarov, Dmitry Konovalov, Alan Herbert, Maria Poptsova

## Abstract

A long-standing question concerns the role of Z-DNA in transcription. Here we use a deep learning approach based on the published DeepZ algorithm that predicts Z-flipons based on DNA sequence, structural properties of nucleotides and omics data. We examined Z-flipons that are conserved between human and mouse genomes after generating whole-genome Z-flipons maps by training DeepZ on ChIP-seq Z-DNA data, then overlapping the results with a common set of omics data features. We revealed similar pattern of transcription factors and histone marks associated with conserved Z-flipons, showing enrichment for transcription regulation coupled with chromatin organization. 15% and 7% of conserved Z-flipons fell in alternative and bidirectional promoters. We found that conserved Z-flipons in CpG-promoters are associated with increased transcription initiation rates. Our findings empower further experimental explorations to examine how the flip to Z-DNA alters the readout of genetic information by facilitating the transition of one epigenetic state to another.

## Introduction

Flipons are important functional elements that regulate various genomic processes by changing conformation ^1–3^. Here we focus on Z-flipons that can form Z-DNA and Z-RNA under physiological conditions. Recent studies highlighted the key-role of Z-RNA in immune response ^4–15^, either switching interferon responses off to limit inflammation or initiating cell death to eliminate virally infected or dysfunctional cells ^14–17^. Other roles for Z-DNA are less well characterized. A number of experiments show that the negative supercoiling produced by RNA polymerases during gene transcription can induce Z-DNA associated with chromatin remodeling by the BRG1 complex ^18,19^. Whole genome experimental approaches for Z-DNA detection are few in number and include human ChIP-seq data sets derived using the Z-DNA structure specific Zα domain ^20^, the use of rapid permanganate/S1 nuclease (Kex) footprinting to detect transient formation of non-B DNA structure in both mouse and human cells ^21^, and kethoxal-assisted single-stranded DNA sequencing (KAS-seq) studies also in mouse and human cells ^22^. Recently a novel Chip-Seq data set for Z-DNA detection in mouse using an anti-Z-DNA antibody in curaxin treated cell was generated ^23^ that was used in this study.

Earlier we developed DeepZ approach ^24^ that makes prediction of Z-DNA forming sites by supervised training on a large number of omics datasets, including histone marks (HM), transcription factor (TF) binding sites, methylations marks, RNA polymerase binding sites, taking into account the *in vitro* determined energetic parameters of dinucleotides switching from B- to Z-conformation ^25^. With DeepZ we were able to generate de novo predictions of Z-DNA forming sequences both in human ^24^ and in mouse ^23^ using experimentally confirmed Z-DNA regions to train and validate the algorithm. With feature importance analysis, we were able to determine epigenetic marks, DNA binding proteins and other factors that are more likely to associate with Z-DNA formation *in vivo*.

Here we reapplied DeepZ approach to human and mouse genome using the common set of omics features. We focused on evolutionary conserved Z-flipons regions between mouse and human as significant regulatory features of these regions should also show conservation, using findings in one genome to validate those from another. We analyzed common and unique associated histone modifications and transcription factors to find those most informative for Z-DNA prediction, then explored different classes of conserved functional Z-flipons located at alternative and bidirectional promoters, and, finally, estimated the effect of Z-flipons on transcription kinetic parameters. We found that conserved Z-flipons are associated with increased transcription initiation rates, have distinct patterns at sites of bidirectional and alternative promoters, and can be clustered into functional groups that reflect known genomic features.

### DeepZ predictions in human and mouse genomes based on the common set of omics features

Here we applied the same DeepZ pipeline as described in ^24^ using human genome ChIP-seq data ^20^ and mouse genome ChIP-seq data ^23^ for training. The schematic of DeepZ approach is presented in Figure 1A. In short, it takes as an input information both from sequence and from available whole-genome experimental maps of various functional elements, i.e., omics data, and additionally it incorporates biophysical information on energy transitions from B- to Z-form.

**Figure 1.**
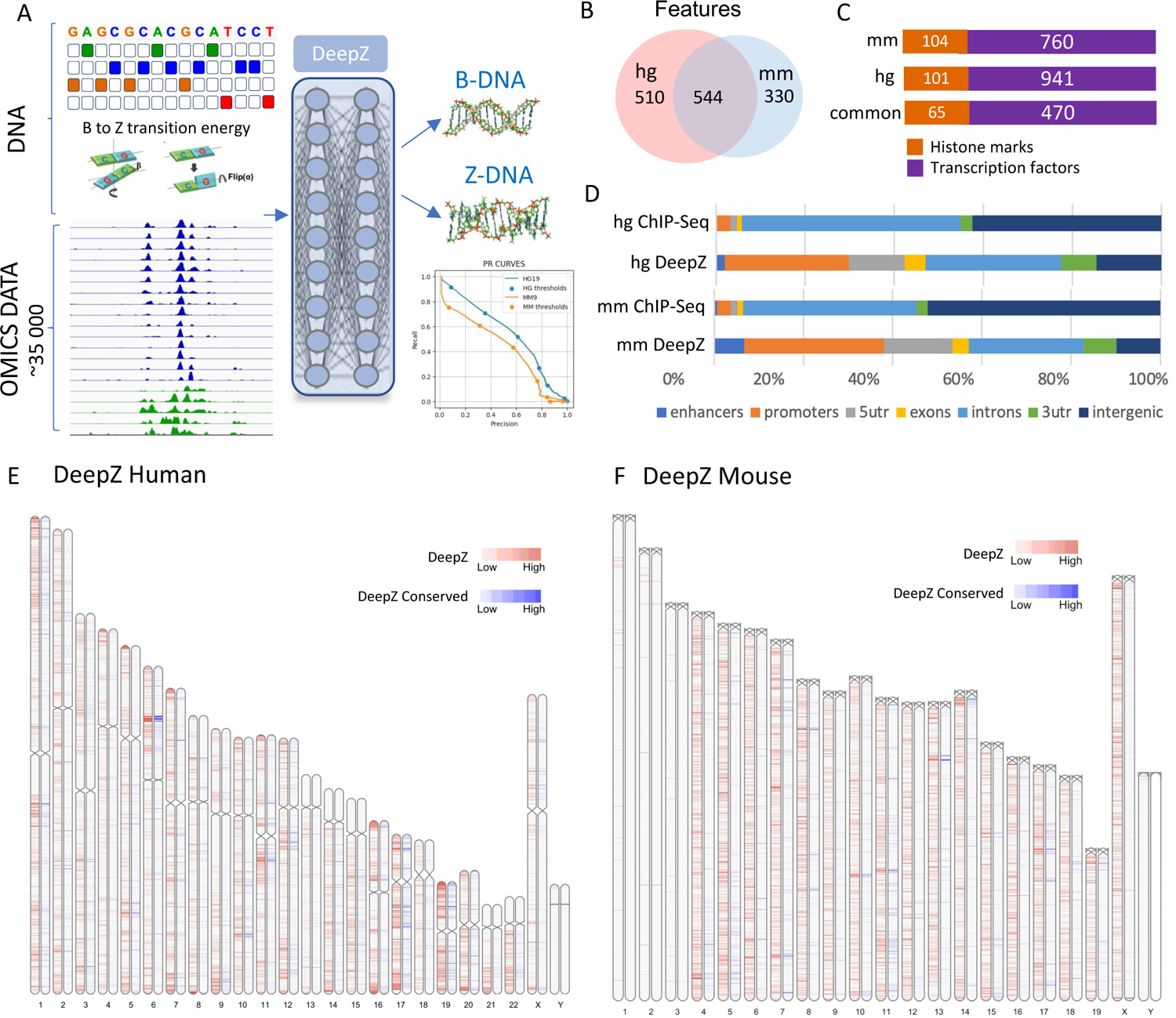
DeepZ predictions in human and mouse genomes based on the common set of omics features. **A.** General Schema of DeepZ approach, with PR curves showing model performance. **B.** Number of common and unique features used in DeepZ model. **C.** Distribution of features over functional groups. **D.** Distribution of Z-DNA regions over genomic regions. **E.** Whole-genome distribution of DeepZ predicted Z-flipons in human genome. **F.** Whole-genome distribution of DeepZ predicted flipons in mouse genome. In **E**, and **F.**, regions conserved between human and mouse are highlighted in blue.

For omics data we took all available histone modifications (HM), transcription factor (TF) binding sites, DNase accessible sites and RNA polymerase binding sites (see full list in Supplementary Table 1). For the purpose of this study, we selected only experiments available for both genomes that resulted in 544 features (Figure 1B-C) including 65 HMs, 466 TF binding sites, 3 methylation maps, 8 RNA polymerase maps, dinucleotide energy transitions from B- to Z-form, and DNase hypersensitivity sites.

We trained the deep neural network as described in ^24^ (see Methods). ROC-, PR-, and F-curves of the DeepZ model performances are presented in Supplementary Figure 1. PR-curve where the precision is low as it is applied to the whole genome without annotation of Z-DNA forming sequences. To overcome this issue, we validated DeepZ by overlap with Z-DANBERT predictions that are based on experimental data that is generalized using a transformer language model algorithm. We present the metrics for 6 different thresholds that are indicated in Figure 1A. For this study we present results for threshold 3 as it has the optimum value for precision and recall as determined using the F1 metric. We generated whole-genome annotations for human and mouse genomes with Z-DNA regions (Supplemental Data 1-2), which comprise 30,083 regions in human and 17,569 in mouse.

The number of potential Z-DNA forming sites in human comprise ∼3 Mb compared to ∼2,6 Mb in mouse genome, but due to the different genome sizes comprise 1% of cumulative genome length in both genomes (Supplementary Table 2). The distributions of the DeepZ predicted Z-DNA over genomic regions for mouse and human are given in Figures 1D. In both genomes the distribution is qualitatively the same with enrichment in promoters, exons, 5’UTR and 3’UTR.

### Conserved Z-flipons are enriched in transcription regulatory factors and active chromatin marks

Next, we investigated how many of the predicted DeepZ regions fall into conserved regions between human and mouse genome (Figure 1E-F, Supplementary Data 3-4). Depending on the quantile threshold (Supplementary Table 2), 11-20% of all predicted Z-DNA regions fell into conserved regions, and the ratio increases up to 24% in human as higher quantiles cut-off are tested. The number of genes with Z-DNA from vertebrate conserved regions in human is almost three times bigger (7188 genes) than in mouse (2310 genes) (Supplementary Table 3). From both lists 966 genes are known human and mouse orthologs that contain predicted Z-DNA sites in the bodies of that genes. GO analysis reveals enrichment of human and mouse orthologs with conserved Z-flipons in regulation of metabolic process (546 genes, FDR e-36), regulation of transcription by RNA polymerase II (220 genes, FDR e-21), response to stimulus (495 genes, FDR e-09), binding to protein and nucleic acids (758 genes; FDR e-21), alternative splicing (614 genes, FDR e-12), chromatin organization (77 genes, FDRe-8), MAPK signaling pathway (37 genes; FDR e-06), with location in nucleus (590 genes, FDR e-45), nuclear lumen (423 genes, FDR e-37) (Supplementary Table 4). Full list of the most-enriched pathways and processes are given in Supplementary Table 5.

### Conserved patterns of transcription factors and histone marks enriched around Z-flipons in human and mouse

We aimed to find common transcription factors and histone marks (here and after referred as an omics feature) that are enriched in regions around conserved Z-flipons. For that we calculated percent overlap of each omics feature with Z-flipons, fold-enrichment of features with Z-flipons over features without Z-flipons and applied permutation randomization test to assess significance (see Methods). We performed this analysis at the genome-wide level, initially examining all regions, then promoter regions, and then CpG- and non-CpG-promoters individually. We selected the top-20 features that have the highest percent of overlap with conserved Z-flipons in each category both for human and mouse and the combined feature importance plot is presented in Figure 2. The full list of TFs and HMs with the values for percent overlap and fold-enrichment for each omics feature and conserved Z-flipons in human and mouse is given in Supplementary Table 6.

**Figure 2.**
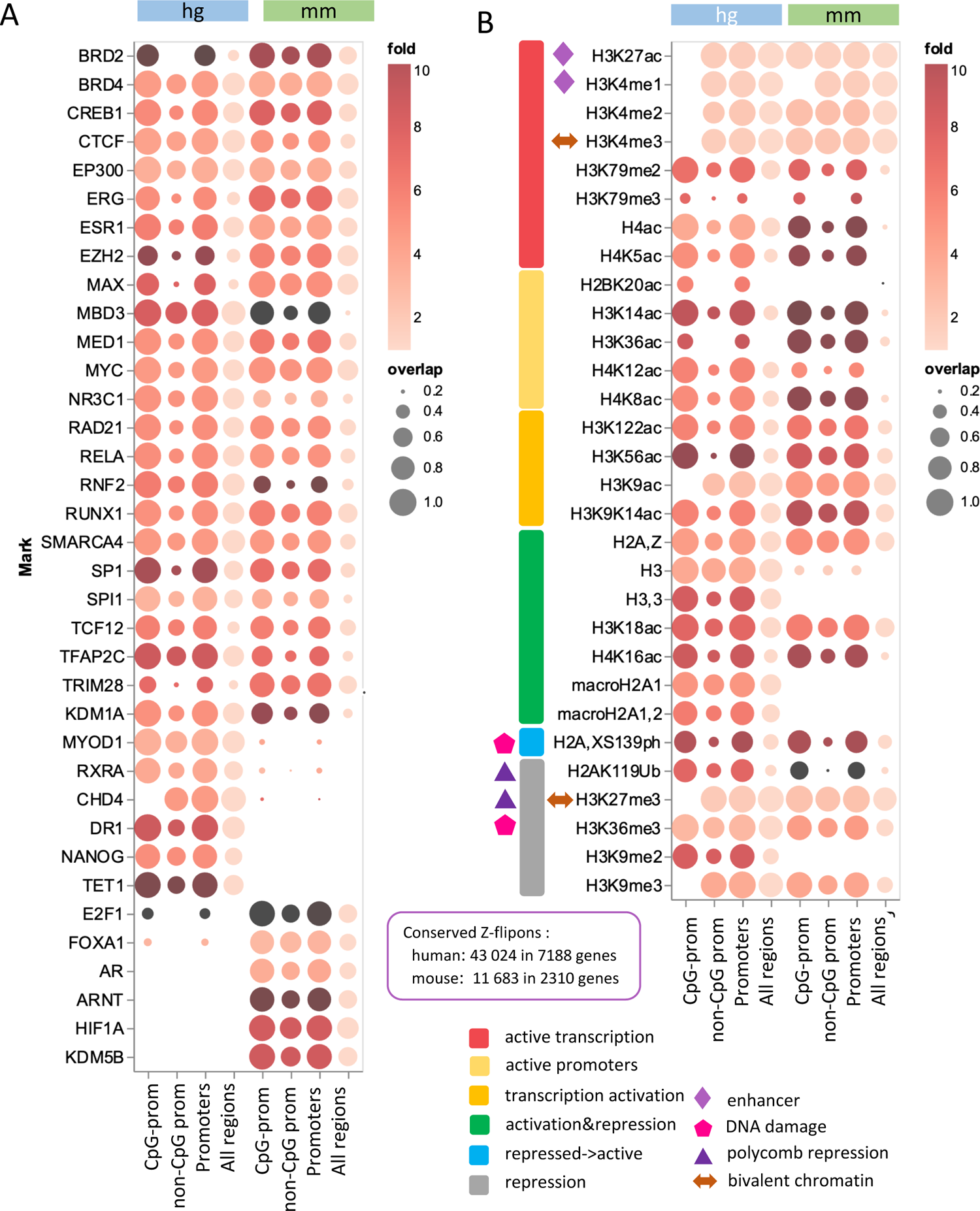
Conserved patterns of transcription factors and histone marks enriched around Z-flipons in human and mouse genome. A. Enrichment of transcription factors around Z-flipons for the entire genome, promoters, CpG-promoters and non-CpG promoters. B. Enrichment of histone marks around Z-flipons.

Figure 2A presents the list of TFs with high percent of overlap with Z-flipons shared between human and mouse genomes. In line with our previous studies ^23,26,27^ we observed enrichment of Z-flipons in promoters. Since GC-repeats are prone to Z-formation we observe significant enrichment of Z-forming sequences in CpG-promoters. However, an enrichment is also observed around non-CpG promoters.

The top TFs that have high colocalization with conserved Z-flipons in promoters in both genomes are BRD2, BRD4, CREB1, CTCF, EP300, ERG, ESR1, EZH2, MAX, and others (Figure 2A). Functional analysis of TFs with more than 90% overlap with Z-flipons in promoters showed enrichment in chromosome organization (FDR e-15), histone modifications (FDR e-9), positive and negative regulation of transcription by RNA Polymerase II (FDR e-23), as well as in developmental process (FDR e-9).

Among histone marks enriched in both genomes (Figure 2B) many are acetylated indicating active promoters and transcription activation. H3K4me1/me2/me3 are also indicators of active transcription. We observe similar enrichment of Z-flipons in marks that are associated both with activation and repression that are usually a hallmark of chromatin state at differentiation. These are histones H2A.Z, macroH2A.1,2, H3.3, H3K18ac, H4K16ac. H3 and H3.3 are associated with chromatin structure. The same analysis for the entire set of DeepZ predicted Z-flipons including non-conserved ones supports that many of the HM and TF features are enriched only in the conserved parts of the genome, highlighting the role of evolutionary selection (Supplementary Figure 2).

### Clusters of common conserved Z-flipons between human and mouse reveal functional groups of LINEs, embryonic development, and neurogenesis

We combined both experimental and DeepZ predicted Z-DNA regions for human and mouse genomes in one data set where each Z-flipon is defined by a vector of omics features. We applied UMAP clustering to this combined human and mouse, predicted and experimental data sets (Figure 3A). We can see that experimental regions are distributed all over the plot with DeepZ predictions expanding the experimental clusters. This reflects the fact that experimentally it is possible to detect only a subset of all available functional Z-flipons, but overall, the number is much larger. For the majority of Z-flipons human and mouse clusters are separated however there exists a common region where both human and mouse omics vectors are close enough (Figure 3B). We extracted coordinates of Z-DNA regions from the common cluster and mapped them to corresponding genes from which we selected orthologs. GO-analysis of orthologs from the common cluster revealed that genes with conserved Z-flipons are enriched in the development and differentiation processes, mainly in neurogenesis, and in particular related to synapse organization and functioning (Supplementary Table 7). Example of Wnt Family Member 5A (WNT5A) human and mouse genes with Z-flipons from the presynapse assembly pathway is given in Supplementary Figure 3. This gene harbors many Z-flipons detected by three methods - DeepZ, experimental KEx permanganate/S1 nuclease detection method,^21^, and also with Z-DNABERT model ^27^. Z-flipons in WNT5A gene are located in 5’UTR, near splice-sites, in exons, at alternative promoters. The one from 5’UTR is composed of GC-repeats and supported by three Z-DNA detection methods.

**Figure 3.**
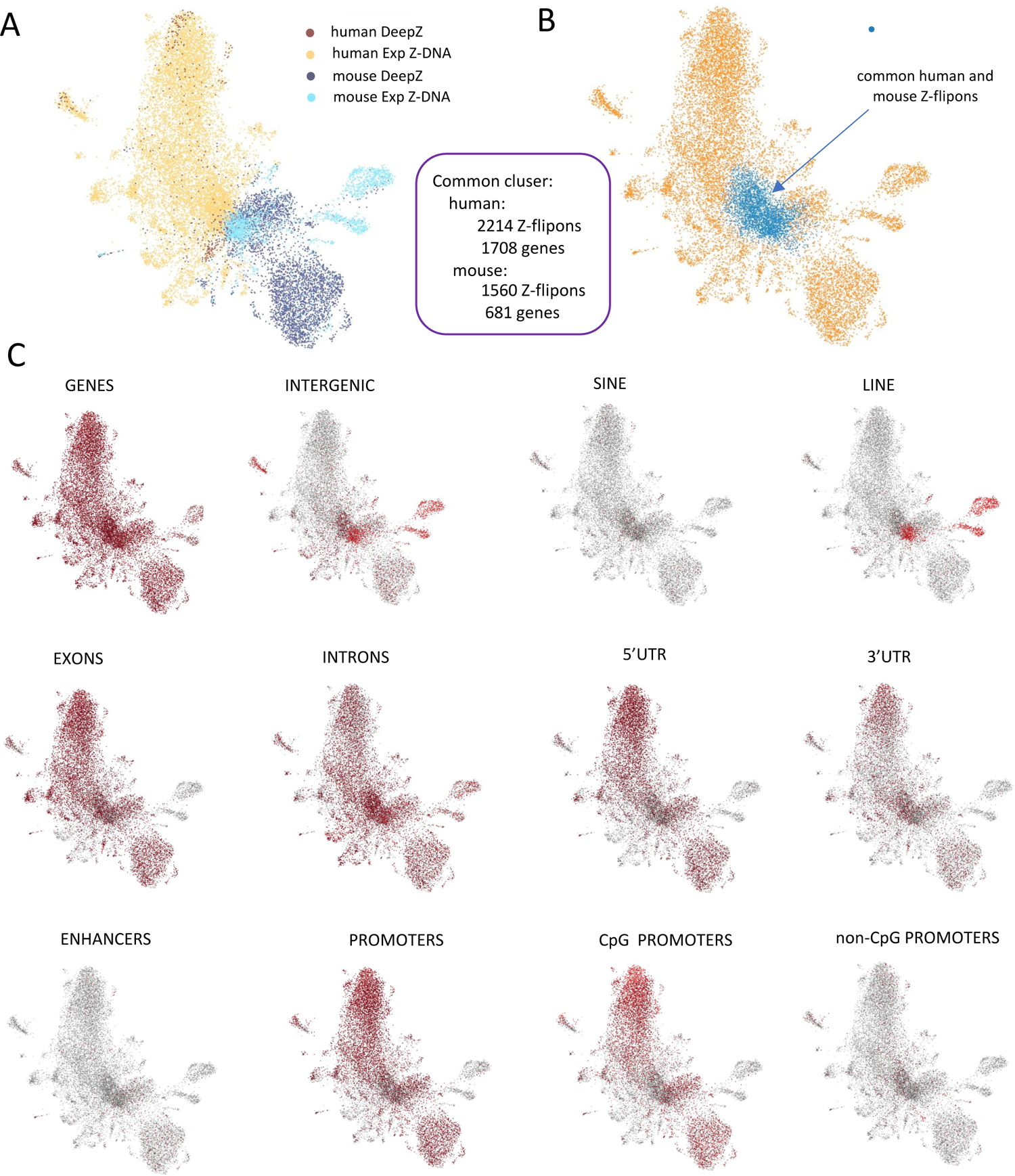
UMAP clustering of Z-flipons based on vectors of common omics features. A. UMAP clustering of experimental and DeepZ predicted Z-flipons in human and mouse genomes. B. Cluster of common human and mouse genome. C. Distribution of Z-flipons over genomic regions, regulatory elements, and transposons.

Figure 3C shows how Z-flipons are distributed over genomic regions. Interestingly, the DeepZ separates the genome by feature, even though such information was not included in the training data. For example, SINE and LINE repeats form a separate cluster in the map. In line with our previous findings, LINES are enriched in curaxin-induced Z-DNA from the mouse genome ^23^ and are enriched for the experimental data (light blue in Figure 3A).

Traditional UMAP clustering revealed functionally similar groups of Z-flipons that are combined together by similar omics feature vectors (Figure 4). Analysis of the top marker features for each cluster revealed that conserved Z-flipons can be assigned to such functional groups such as embryonic development and morphogenesis, RNA Polymerase II, RNA Polymerase III, heterochromatin LINEs, chromatin binding, negative regulation of transcription, cellular component organization.

**Figure 4.**
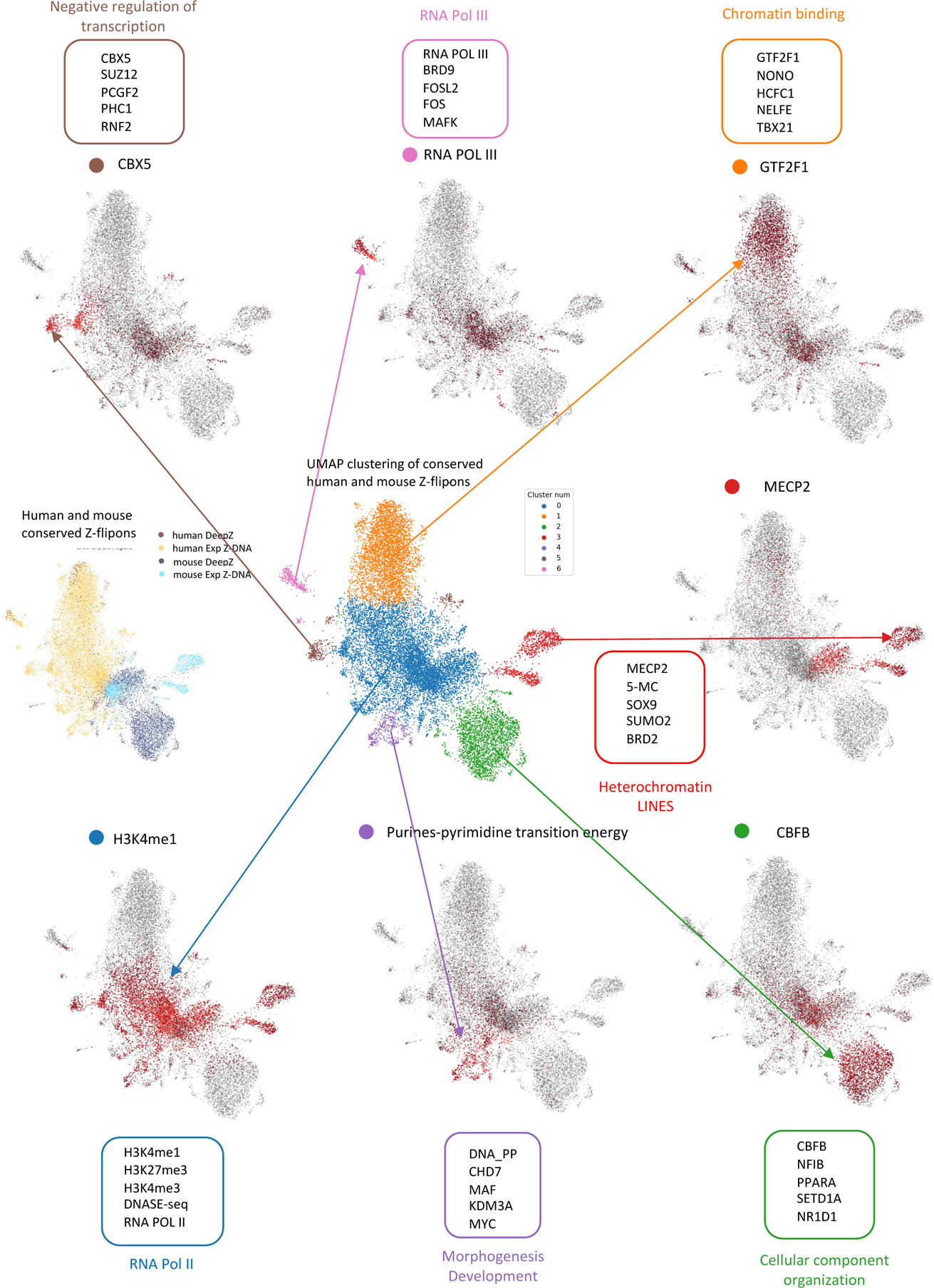
UMAP сlusters of human and mouse Z-flipons with one top marker feature highlighted for each cluster. The first 5 marker features are given in the boxes.

This analysis independently confirmed that Z-flipons have conserved functional roles in both genomes that revealed by common associated defining features from omics vectors (for all important markers in UMAP clustering see Supplementary Table 8).

### Functional Z-flipons at promoters

We found that 15% and 7% of conserved Z-flipons fell in alternative and bidirectional promoters. Below we explore in more detail cases when DeepZ predicted Z-DNA region was detected in both human and mouse orthologs by two other methods that are orthogonal to the DeepZ approach i.e., KEx and Z-DNABERT.

### Z-flipons at alternative promoters

Many genes have alternative promoters but the mechanism of activation and how they control gene expression is not clear. Our analysis revealed that, depending on the size of the upstream region, 10-15% of DeepZ-predicted Z-flipons overlap with alternative promoters (Supplementary Table 9). Here we highlight different cases with alternative promoters are very close to transcription start site or located at a distance from the main promoter. Plekha7 (Pleckstrin homology domain-containing family A member 7) is one of the conserved gene from the major cytoplasmic scaffolding and adaptor proteins of vertebrate apical junctions ^28^ and is one of the clusters of Plekha genes that originates early in chordate evolution ^29^. PLEKHA7 signaling is necessary for the growth of mutant KRAS driven colorectal cancer ^30^ with extensive alternative splicing towards the 3’end of the gene (Figure 5A-B). The alternative PLEKHA7 promoters overlap with the Z-flipons detected by three different methods (DeepZ, KEx and ZDNABERT) (Figure 5 C,D). The alternative promoters in both genomes have noticeable column of ChIP-seq signals for HMs and TFs detected as enriched in Z-flipon regions in both genomes and aggregated for all tissue types (Figure 5 C,D). Because DeepZ was trained on a broad ChIP-seq data, the width of DeepZ prediction is comparable to ChIP-seq peak widths and does not have the higher resolution possible with KEx and Z-DNABERT. Human Z-flipon for distant alternative promoter (left column at Figures 5 C,D) is composed of CA-repeats (GT-repeats at the opposite strand), while mouse Z-flipon does not have this pattern. The right column of omics features at the main promoter (right column in Figure 5 C,D) also has Z-flipons confirmed by three methods. Splicing graph (Figure 5 A,B) shows that there are alternative promoters nearby start sites for this gene. DeepZ predicted two Z-flipons around this promoter based on two peaks of omics features corresponding to tandem promoters. Since DeepZ were trained on aggregated experimental signals from different tissues, we explore how omics signal are distributed in individual tissue type. Here we show examples of feature distribution for liver in human and mouse genome (Figure 5 E,F). The H3 methylation pattern is consistent with the absence of expression in liver.

**Figure 5.**
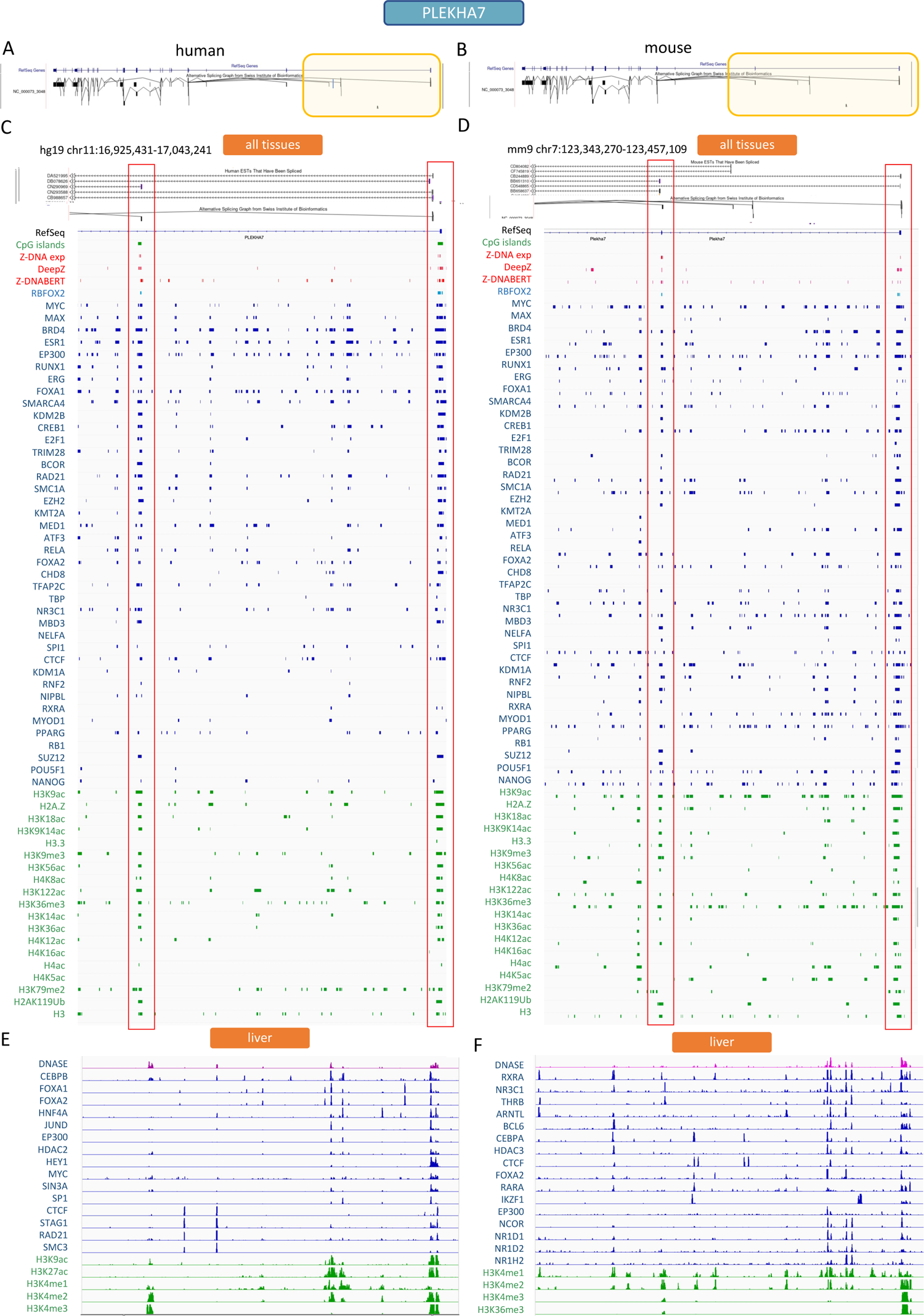
Z-flipons at alternative promoters of PLEKHA7 orthologs in human and mouse genome. **A-B.** Alternative splicing graph for PLEKHA7 in (**A**) human and (**B**) mouse genomes. **C-D.** Region of PLEKHA7 in human (**C**) and mouse (**D**) genomes with Z-flipon signals detected by three methods – DeepZ, KEx, Z-DNABERT, and signals from omics features enriched in Z-flipons both in human and mouse genomes. Omics features are aggregated signals from all tissues as they were used in DeepZ model. **E-F.** Selected omics features for human and mouse genome for liver tissue type.

### Z-flipons at bidirectional promoters

Our analysis revealed that 4-7% of DeepZ-predicted Z-flipons overlap with bidirectional promoters. We observed cases where Z-flipons were detected with three methods (KEx, DeepZ, Z-DNABERT) at bidirectional promoters (Supplementary Table 9). Example of Z-flipons in human and mouse genome at the bidirectional promoter of transmembrane protein 51 TMEM51 (transmembrane protein 51) and long non-coding RNA (lncRNA) TMEM51-AS1 (TMEM51 Antisense RNA 1) are presented in Figure 6. Aggregated omics signal span a region covering both bidirectional and alternative promoters. Many signals are split in two, three, or even four separated peaks showing differential use of promoters between tissues. This can be clearly seen in tracks for specific tissue type. In blood, there is differential binding of TF to each promoter region. TFs with two peaks – CTCF, RNF2, SUZ12, RUNX, ERG, and with three peaks – FOXA1, FOXA2, CREB1,RAD21 (see Figure 6) are those involved in chromatin binding and regulation of transcription by RNA pol II. There is a Z-flipon in intron confirmed also experimentally that consists of GT-repeats (Figure 6 A,B). Since it is not located at promoter, its function can be related to alterative splicing (see the example of MORN1 below). The lack of TMEM51 expression in blood is is consistent with the H3 methylation marks present in both human and mouse(Figure 6 F,H).

**Figure 6.**
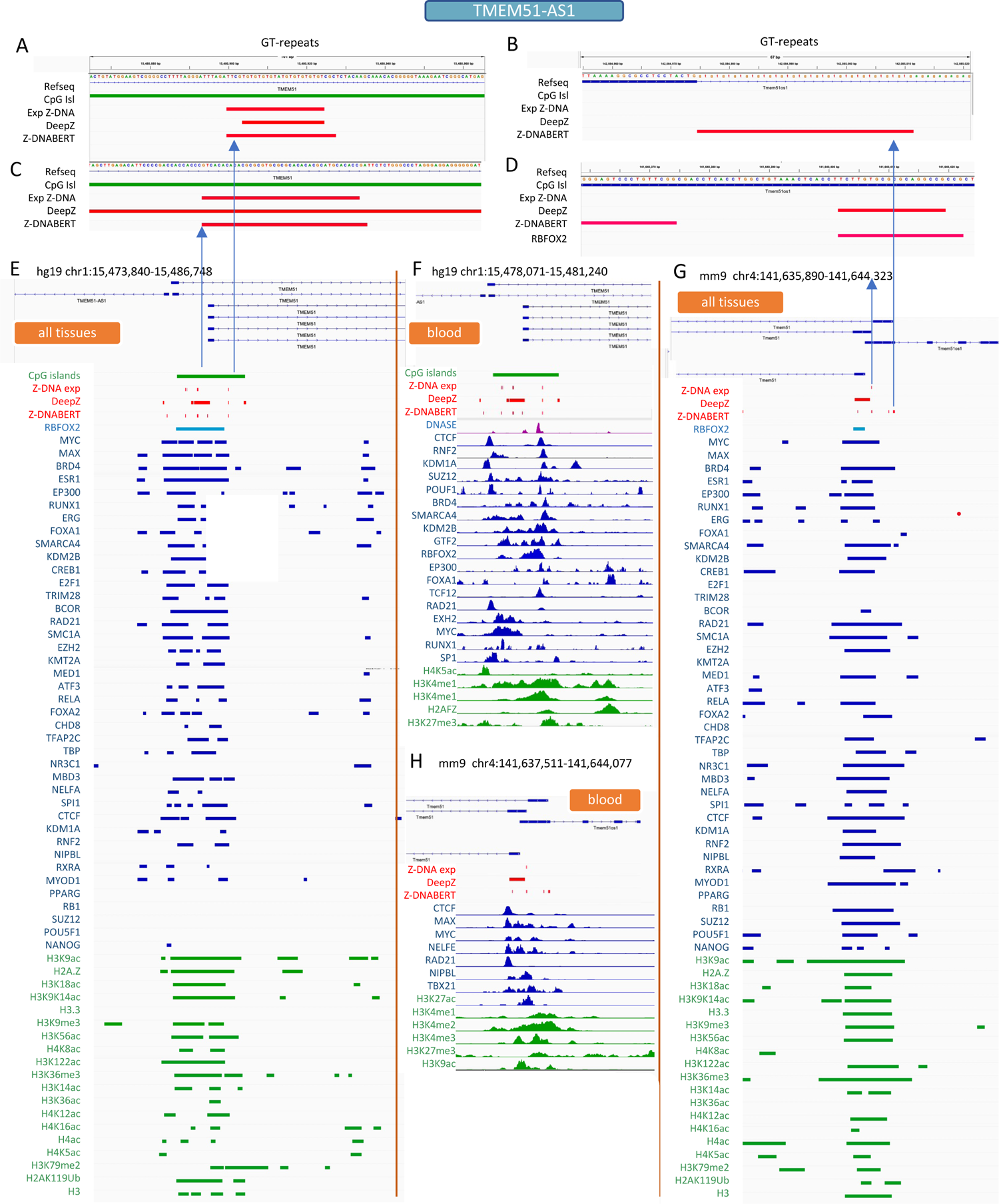
Z-flipons at bidirectional and alternative promoters of TMEM51-AS in human and mouse genomes. **A-D.** Enlarged regions around Z-flipons detected by three methods – DeepZ, KEx, Z-DNABERT in human (**A,C**) and mouse (**B,D**). **E-G.** Region around bidirectional promoter of TMEM51-AS and signals from omics features enriched in Z-flipons both in human and mouse genomes. Omics features are aggregated signals from all tissues as they were used in DeepZ model (**E,G**). Selected omics features for human and mouse genome for blood tissue type (**F**).

The same conserved multiple peaks pattern for omics signal is shown for BRCA1-NBR2 bidirectionally transcribed pairs (Supplementary Figure 4) with Z-flipons at bidirectional and alternative promoters in both genomes. For BRCA1-NBR2 pair we can also see that bidirectional promoter is located close to alternative promoters, and omics signals have two peaks, especially HMs, as it is more clearly seen at tracks for liver tissue (Supplementary Figure 4 C,D). Z-flipons in intron of NBR2 is also composed from repeats similar to intronic Z-flipons in TMEM51-AS1. The pattern of omics code for bidirectional gene pairs is complex and evolved in a way so that it has to separate usage of bidirectional and alternative promoters.

### Association of Z-flipons on transcription initiation rate

We explored the association of Z-flipons on transcription kinetics parameters with the data from ^31^. We found significant difference in distributions of initiation frequencies for promoters with conserved DeepZ flipons as compared to promoters without Z-flipons (Figure 7A) where initiation frequency is higher with conserved Z-flipon promoters. We also see the difference in distributions for CpG-promoters with conserved Z-flipons (Figure 7B). We also see the significant difference between CpG- and non-CpG promoters even without Z-flipons (Figure 7 C). There was no difference in pause time of elongation rate associated with Z-flipons.

**Figure 7.**
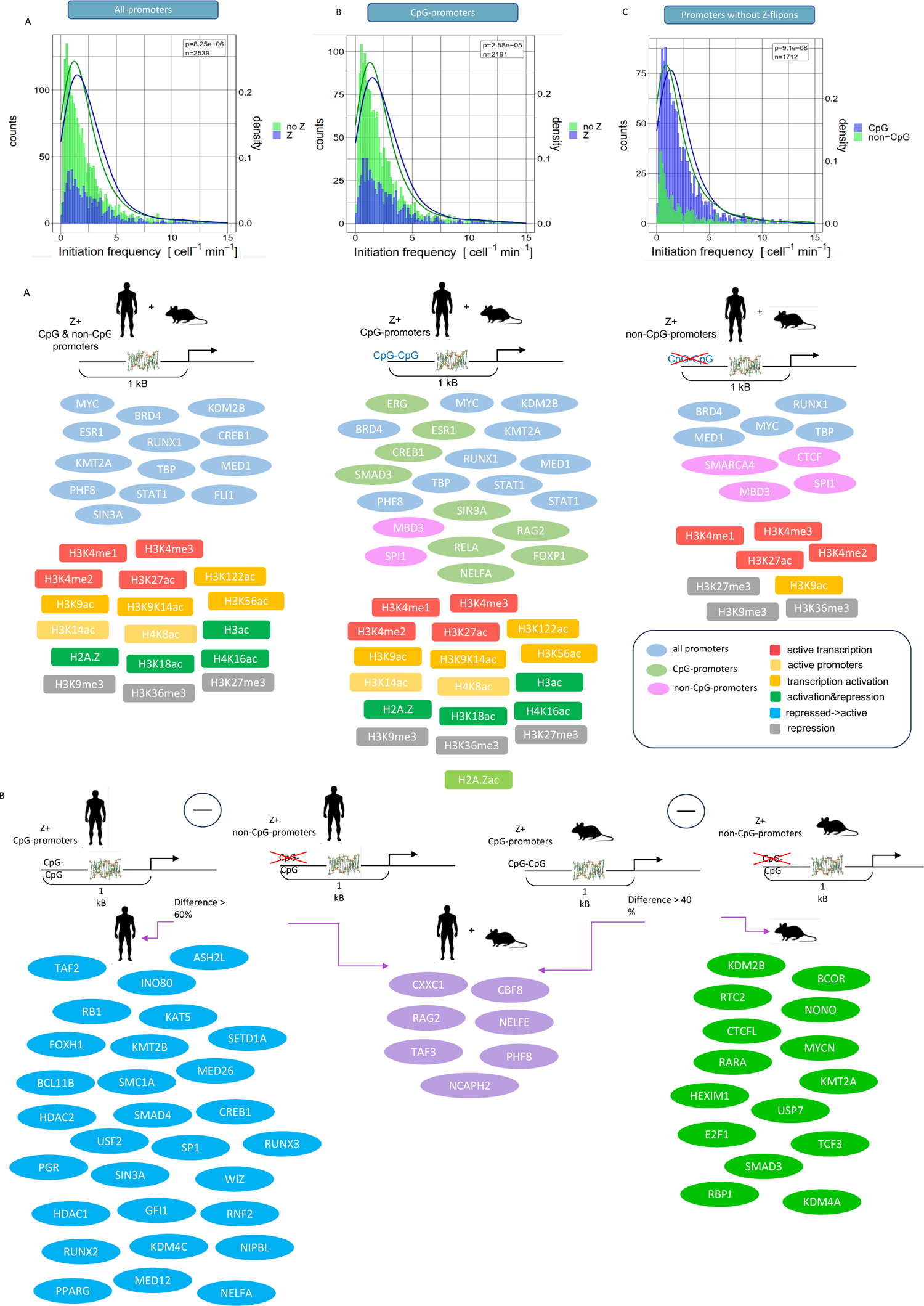
**A-B.** Transcription initiation rate for different types of promoters with or without Z-flipons. **A.** Transcription initiation rate for promoters with conserved Z-flipons is larger than the rate for promoters without Z-flipons. **B.** Transcription initiation rate for CpG-promoters with conserved Z-flipons is larger than rate for CpG-promoters without Z-flipons. **C.** Transcription initiation rate for CpG-promoters with Z-flipons is larger than the rate for non-CpG promoters with Z-flipons. **D-E**. Common and unique omics features for CpG and non-CpG promoters with conserved Z-flipons. **D**. Omics features enriched around all, CpG- and non-CpG promoters. **E**. Transcription factors that show the difference more than 60% in human and more than 40% in mouse between CpG- and non-CpG promoters with Z-flipons. TFs that are more characteristic for CpG promoters in human are depicted in blue, for mouse – in green, common for human and mouse – in lilac.

The summary of how most enriched TFs and HMs are distributed over CpG and non-CpG promoters are given in (Figure 7D). CTCF, SMARCA4, MBD3, SPI1 (chromatin organization and negative regulation of transcription by RNA Polymerase II) are more associated with non-CpG promoters. ERG, ESR1, CREB1, SMAD3, SIN3A, RAG2, RELA, NELFA, FOXP1 are all involved in regulation through chromatin remodeling, cell growth and differentiation are associated with CpG promoters. The top TFs for which the largest difference in overlap between CpG- and non-CpG promoters and are common for human and mouse are methylated histone binders (CXXC1, PHF8, RAG2), chromatin binders (NELFE, NCAPH2), and have zinc finger domain (TAF3, PHF8, RAG2, CXXC1) (Figure 7E). Many of these factors may have direct or indirect influence on transcription kinetic parameters.

### Z-flipons and exon skipping

MORN1 (MORN repeat containing protein 1) has Z-flipons detected by three methods (KEx, DeepZ and ZDNABERT) in human and mouse genome in the intron near the exon that can be skipped (Figure 8). Examination of the Z-DNA sequence showed that it consists of GT-repeats (CA-repeats at the reverse strand). The set of histone marks and transcription factors overlapping this flipon is not that large as for the alternative or bidirectional promoters (Figure 5-6). The peaks are seen for MYC, BRD4, ESR1, RUNX, ERG, FOXA1, SPI1, CTCF – all are involved in chromatin organization and remodeling. Histone mark signals are seen for H3K9me3, H2A.Z, H3K18ac, H4K14ac, H3 – marks of both active and repressive chromatin characteristic for developmental processes. In both genomes Z-flipon forming region consists of GT-repeats, however in mouse the region is 440 bp long while in human it is about 50% smaller and distorted by mutations. Chromatin structure is likely to play a role in alternative splicing as well as the secondary structure formed by the transcript. The role Z-DNA and Z-RNA play in these outcomes is currently underexplored.

**Figure 8.**
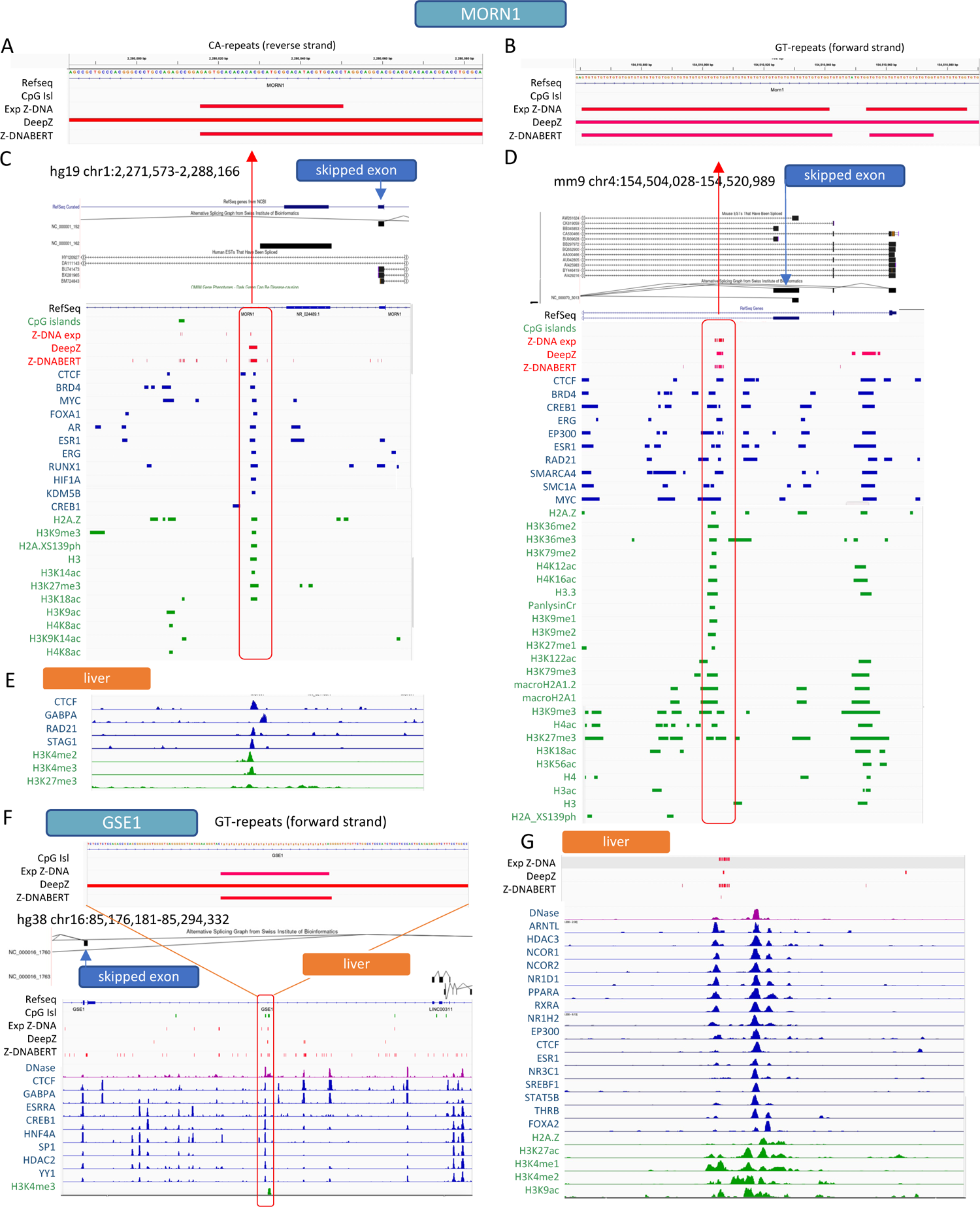
Z-flipons formed at GT-repeats and exon skipping. Example of MORN1 (**A-E,G**) and GSE1 (**F**) genes. Enlarged regions depicting Z-flipons composed of GT-repeats and detected by three methods – DeepZ, KEx, Z-DNABERT in human (**A**) and mouse (**B**). **C,D**. Region around Z-flipon in intron of MORN1 gene and signals from omics features enriched in Z-flipons both in human (**C**) and mouse (**D**) genomes. Omics features are aggregated signals from all tissues as they were used in DeepZ model (**E,G**). Selected omics features for human (**E**) and mouse (**G**) genome for blood tissue type. **F**. Z-flipons formed at GT-repeats in intron of GSE1 gene in human genome. Z-flipon is detected by three methods – DeepZ, KEx, Z-DNABERT. Selected omics features are presented for liver tissue type.s

## Discussion

Here we found evidence that conserved Z-flipons are associated with increased transcription initiation rates. We checked the transcription kinetic parameters for CpG and non-CpG promoters with and without Z-flipons and found significant difference in distributions specifically for CpG promoters with conserved Z-flipons and without any flipons. This is in line with the mechanism of transcription regulation by resetting promoters with Z-flipons as proposed in ^32^ and with the regulation of TATAA box promoters by sequence-specific transcription factors that give a broader dynamic range of expression than is possible by chromatin modification of histone alone. Here the energy of negative supercoiling can be used to rapidly displace the promoter initiation complex where binding of components positively supercoils DNA to open up the transcription bubble. Other transcription kinetic parameters such as polymerase pause duration and transcription elongation rates can be further tuned by micro-RNA and non-B flipons. Polymerase pauses 100-150 bp downstream of transcription start sites where Z- or G-flipons can be located.

We presented an analysis of conserved Z-DNA regions in human and mouse genomes predicted by the deep learning model DeepZ. The model was trained using experimentally confirmed Z-DNA regions and incorporates information both from sequence, energetic parameters related to Z-DNA formation, and omics data from a wide range of ENCODE ChIP-seq experiments. The analysis allowed us to determine conserved patterns of colocalization of TFs and HMs that colocalize with Z-DNA regions. In general, Z-flipon in conserved promoters and gene bodies are associated with factors that direct chromatin remodeling and promote higher rates of transcriptional initiation.

From more than 500 common TFs and HMs used to train the DeepZ model we selected top-20 from the list of TFs and HMs and explored their patterns at the genome-wide level and at the promoters, both CpG and non-CpG. Features with more than 90% overlap with Z-flipons in promoter regions both in human and mouse include many transcription factors participating in chromatin organization (BRD2, BRD4, MBD3, EZH2, CTCF, EP300, ESR1, MYC, NR3C1, RAD21, RELA, SMARCA4, SPI1(FDR 6.07e-10)); chromatin remodeling (BRD2, BRD4, MBD3, MYC, ESR1, EZH2, SMARCA4 (FRD 5.14e-6)); positive regulation of miRNA transcription (MYC, NR3C1, SPI1, RELA, SMARCA4 (FDR 7.17 e07)); epigenetic regulation of gene expression (SpI1, CTCF, EP300, EZH2, MBD3 (FDR 4.89e-5)) (see Figure 2 and Supplementary Table 6). Colocalization of these transcription factors with Z-flipons are conserved at the promoter level. Conserved Z-flipons at non-CpG promoters are characterized by SWI/SNF superfamily-type complex (SMARCA4, CTCF, MBD3) coupled with RNA polymerase II transcription regulator complex (BRD4, MED1, MYC, RUNX1,TBP).

Expanding analysis to the non-conserved Z-flipons, we can see that the pattern of histone marks at CpG-promoters are similar in human and mouse however the difference is apparent at non-CpG promoters (Supplementary Figure 2). CpG-promoters with Z-flipons contain many acetylated marks (H3K18ac, H4K8ac, H3K14ac, H3K36ac, H4K12ac, H2BK20ac) and show high enrichment in both genomes. Non-CpG promoters with Z-flipons are lacking most of the histone marks except for two enhancer marks H3K27ac and H3K4me1, and the marker of bivalent chromatin H3K4me3. This suggests that regulation of non-CpG promoters is implemented utilizing some other mechanism rather than via histone modifications, and other types of flipons (non-B structures) alone or coupled with miRNA play a significant role ^32,33^. In numerous cases alternative transcription start sites are located at a distance of 10-50 bp, which is one third or less of the DNA region wrapped around a nucleosome containing a histone mark. Z-flipons can help to find the precise localization of alternative TSS. Z-DNA can also play a role in preventing transcriptional interference by preventing an RNA polymerase from an upstream promoter from disrupting the transcription complex formed at a downstream TSS ^32^.

Histone patterns are also similar at human and mouse functional Z-flipons. H2A.Z overlap more than half (63% and 66%) of DeepZ conserved regions in mouse and human. Almost all CpG-promoters with Z-flipons (97% in human and 99% in mouse) are marked with H2A.Z which interestingly plays a role in neuronal activity and differentiation ^34,35^ and has a lysine/alanine dipeptide repeat that can dock to Z-DNA ^36^. The acetylation of this variant H2A.Zac is higher in mouse compared to human (94% vs 77% at CpG-promoters); it activates neo-enhancers in prostate cancer ^37,38^ and plays a role in trophoblast differentiation in human embryonic stem cells ^39^. Acetylation of histone H3 regulates developmental program in early human embryos ^40^. H3 acetylation is dynamic and varies during preimplantation development window. CpG-promoters with Z-flipons have 93% overlap with H3ac in human and 95% in mouse. H3K122ac (86% and 95% overlap at CpG-promoters in human and in mouse) is a coactivator p300/CBP and induce perturbation of nucleosomes ^41^ and along with H3K16ac, H3K18ac and H3K56ac disrupt chromatin structures and are target of sirtuin deacetylates that suppress gene expression ^42^. Interestingly, H3K18ac (96% and 99% at CpG-promoters in human and mouse) primes mesoderm differentiation ^43^, and this mark was proposed to be used as a marker of cancer progression and potential target of anti-cancer therapy ^44^. DNA demethylases can recognize H3K18ac mark and then be recruited to the chromatin. HDA6 could erase H3K18ac mark to prevent DNA demethylation ^45^. Indeed, in human CpG promoters H3K3me1,2, and 3, H3K27me3 and H3K9me3 are diminished. The role of Z-flipons in particular and other types of flipons in general in chromatin organization is underestimated. Earlier we developed deep learning model that can recognize flipons as nucleosome barriers ^46^ however how flipons add to chromatin remodeling is still to be investigated.

UMAP analysis of close omics feature vectors defining Z-flipons in more than 544-dimensional space revealed common functional clusters shared between human and mouse conserved Z-flipons. These clusters overlap with genomic regions and transposons, specifically with LINEs in mouse data. This confirms our previous finding of enrichment of LINEs in curaxin-induced data ^23^. SINEs are routinely cleaned out by the processing pipeline used for processing experimental data and are depleted in our training sets, likely explaining why we do not observe enrichment of Alus in our analysis.

The functional groups identified with a standard marker analysis can be divided into common and human- and mouse-specific. A big cluster covering both human and mouse flipons is associated with enhancer mark H3K4me1 and RNA POLII and DNase accessibility sites. The POLII general transcription factor (GTF2F1) that has promoter-specific chromatin binding activity marks a chromatin binding cluster of human Z-flipons. CBFB (core binding factor beta), a master regulator of genes specific to hematopoiesis and osteogenesis, marks another large cluster of mouse-specific cellular component organization. MECP2 (methyl-CpG binding protein 2), a reader of DNA methylation, marks a cluster, that is also a LINE cluster in mouse genome. Z-flipons with a strict purine-pyrimidine alternation were clustered together with morphogenesis related genes that are evolutionarily old. This alignment suggests the early evolutionary selection of Z-flipons composed of GT- or GC-repeats. CBX5 (chromobox 5) marks a small cluster of heterochromatin forming factors that participate in negative regulation of transcription by binding to methylated histone lysines. There is a small cluster of RNA Pol III associated Z-flipons that reflects the abundance of Z-DNA forming potential in ribosomal DNA. All these clusters include common human-mouse conserved set of Z-flipons.

As it was shown by us earlier Z-flipons are enriched in promoters. Enrichment in promoters of DeepZ conserved human-mouse Z-flipons is even higher. We found common pattern of omics features colocalized with conserved Z-flipons at bidirectional promoters. Example of TMEM51 gene bidirectionally transcribed with long non-coding TMEM51-AS1 represents a pattern of TFs and HMs characteristic for alternative promoters, specifically and for tandem promoters when an upstream promoter suppresses downstream ^47^. Long non-coding RNA TMEM51-AS1 is differentially expressed in cancer ^48–50^.

Around 52% of human RefSeq genes are subject to regulation by alternative promoters^51^. They can be located in the first 10-50 bp of the main promoter or at a distance of more than 1 Mb. The distant alternative promoters are marked with the same pattern of omics features as the major one, however there is no place to mark close alternative promoters, and here Z-flipons (and other types of flipons) can play a role of a separator and a switch ^32^.

## Conclusion

Here we present evidence for association of conserved Z-flipons in CpG-promoters with transcription imitation rates that is in line with a mechanistic function of Z-flipons in resetting of transcription initiation complex. We investigated the properties of conserved human and mouse Z-flipons associated with common set of TFs and HMs and revealed common patterns of omics code defining conserved Z-flipons at bidirectional and alternative promoters. We revealed existence of TF and HM associated with Z-flipons that are conserved during the evolution of both human and mouse. The strict purine-pyrimidine repeats that favor Z-DNA formation are still under the selection and participate in developmental processes and morphogenesis. In other regions of the genome, DeepZ is able to identify differences in genomic regions that differ in their functional roles based only on the prediction of Z-flipons at these locations rather than being explicitly trained to do so.

## Methods

### Deep Z approach

We applied here DeepZ approach as described in detail in ^24^. As input data DeepZ accepts matrices of size L x W, where L equals to 5000 bp and W represents the number of features from omics data. Input features consist of two major groups: the first group incorporates information on DNA sequence, it includes one-hot-encoded DNA sequence of length L, boolean features representing different simple repeats, and energy transitions to switch from B- to Z-conformation for each dinucleotide as it is implemented in ZHunt ^25,52^. The second group of features comprise information from omics data and includes histone marks (HM), DNase I hypersensitive sites (DNase-Seq), transcription factors (TF) and RNA-polymerase (RNAP). The omics data is taken from chip-atlas.org. Since different sets of experiments available for human and mouse genomes, we took intersection of features that are present in both genomes (Figure 1C). The distribution of feature groups can be found in Figure 1E. Each feature is linearly scaled to the interval [0, 1]. Full list of features is given in Supplementary Table 1. As a training set we took ChiP-seq data from Zhang et al. ^23^ for mouse genome and ChIP-seq data from Shin et al. ^20^ for human genome.

For experimental Z-DNA ChIP-seq data, both mouse and human genomes were divided into equal segments of the length L. Segments from blacklisted regions ^53^ were excluded. Similarly, segments with more than a half of the nucleotides undefined in the genomic assembly were also excluded. All the segments were marked based on the presence of Z-DNA sites as 1 and 0. All Z-DNA regions were assigned as positive class. The negative class was composed from non-Z-DNA segments twenty-fold in size. The resulting segments were stratified into five folds where 4 folds were included in a training set, and the remaining fold formed a test set. We took the best RNN-based model architecture as in ^24^. All training parameters could be found in source code.

### Whole-genome annotation with Z-DNA regions

The trained DeepZ model was used to generate Z-DNA whole-genome annotation. The model predicted probability to form Z-DNA for every segment from all five folds. We generated the annotation with seven thresholds where i-th threshold equals to the quantile 1 - 10^i.

### Conserved Z-DNA regions

Conserved Z-flipons were identified by overlapping DeepZ predicted Z-flipons with vertebrate conservation tracks from UCSC genome browser: Vertebrate Multiz Alignment & Conservation (100 Species) for human and Vertebrate Multiz Alignment & Conservation (60 Species) for mouse.

### Enrichment analysis for omics features

Association of Z-flipons with transcription factors or histone marks was calculated using Monte-Carlo method (n=1000). Promoter regions were defined as regions of 1000 bp upstream and 200 bp downstream of TSS. TSS locations were based on Gencode v44 ^54^ annotation. UCSC CpG islands ^55^ (CpGI) track was used for CpG-/nonCpG-promoter classification. Promoter was considered CpG if it had an overlap with the CpGI (CpG Island) annotation.

### Transcription kinetics parameters analysis

For each transcript from ^31^ CpG/non-CpG label was assigned in a way described above. P-values were calculated for 6 different DeepZ thresholds (all predictions and conservative) using KS-test. After that they were adjusted for multiple testing (FDR). Initiation rate plots were generated via custom R scripts.

### UMAP clustering of Z-flipons based on omics vectors

In our study, we aimed to visualize all predicted Z-DNA regions using a UMAP two-dimensional embedding. For each interval, we extracted the omics representation and then computed the intensity (number of non-zero base pairs in every interval) of every feature within these intervals. Subsequently, we normalized the mean of each feature across each organism. The vectors derived from these calculations were utilized as feature vectors for the subsequent UMAP analysis. For the UMAP projection, we employed the MANHATTAN metric and selected 55 nearest neighbors to ensure a connected representation. All other parameters were set to their default values.

For segmenting the Z-DNA regions represented in the UMAP, we applied the Leiden clustering method based on signal intensities. A resolution parameter of 0.25 was chosen for this purpose. The same approach was used to identify the central cluster in the UMAP, but with a higher resolution setting of 1.

To identify distinguishing features of a cluster, we compared the mean and variance of every feature both within and outside each cluster, utilizing Z-statistics. A high Z-statistic indicated that a feature was representative of that particular cluster.

