## Supplementary Figures for "Z-Flipons conserved between human and mouse are associated with increased transcription initiation rates"

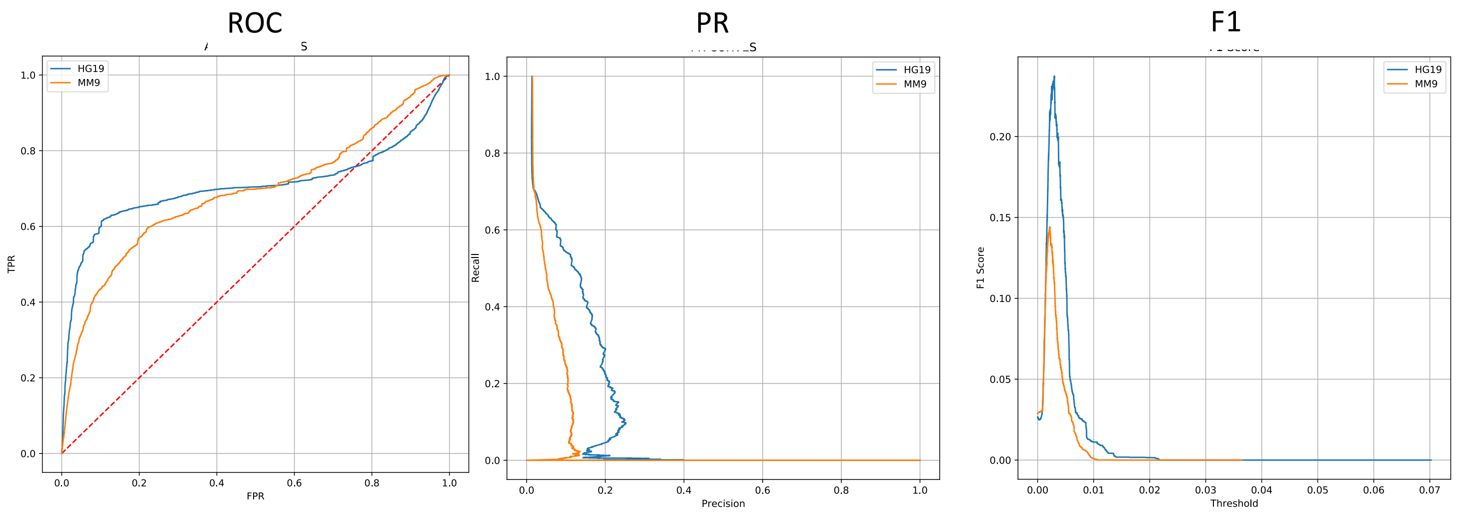


**Supplementary Figure 1.** ROC-, PR-, and F-curves of DeepZ models for human and mouse genomes against each genome without any annotation for flipons. DeepZ models were trained using the same number of omics features.

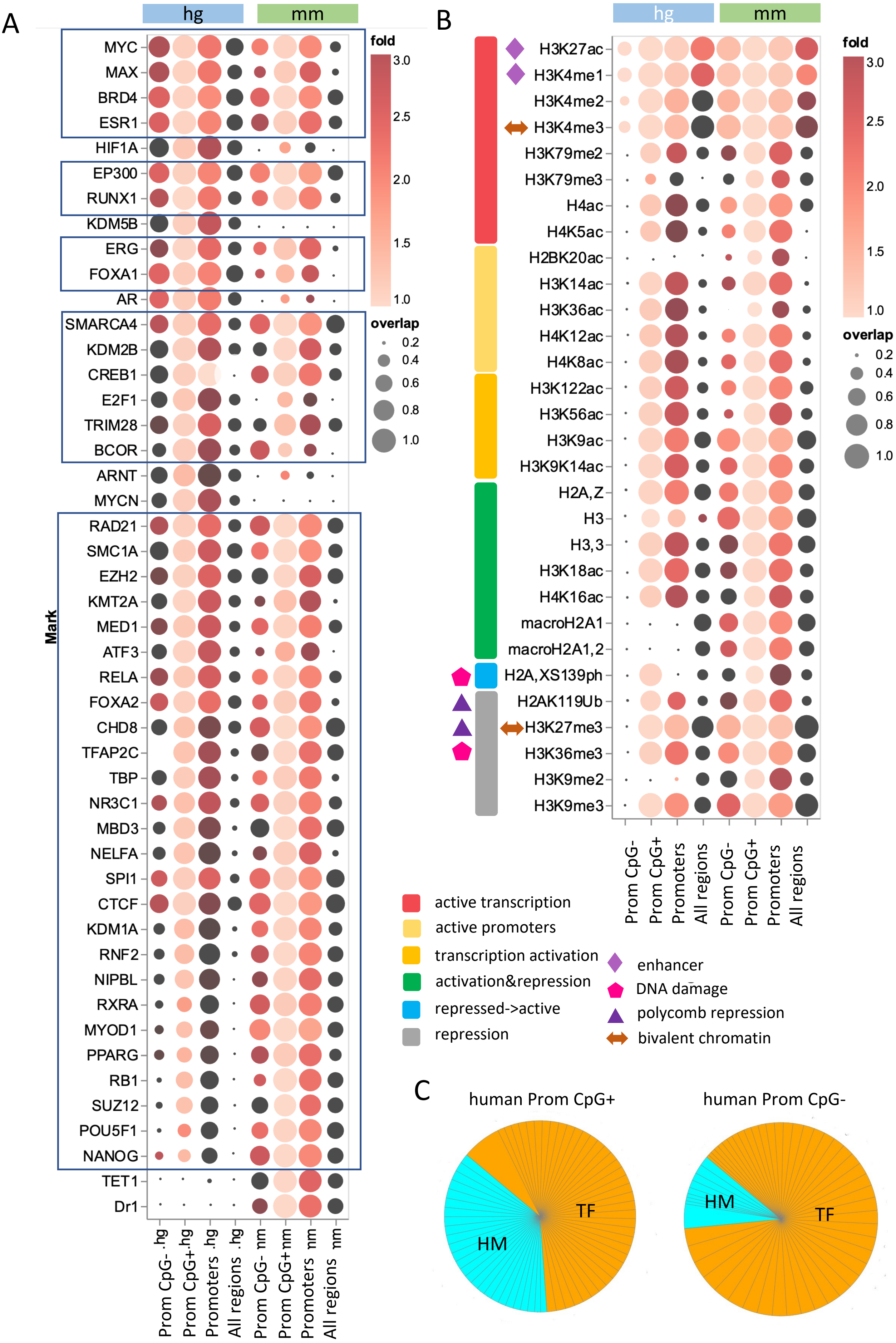
A
hg
mm
MYC
MAX
BRD4
ESR1
HIF1A
EP300
RUNX1
KDM5B
ERG
FOXA1
AR
SMARCA4
KDM2B
CREB1
E2F1
TRIM28
BCOR
ARNT
MYCN
RAD21
SMC1A
EZH2
KMT2A
MED1
ATF3
RELA
FOXA2
CHD8
TFAP2C
TBP
NR3C1
MBD3
NELFA
SPI1
CTCF
KDM1A
RNF2
NIPBL
RXRA
MYOD1
PPARG
RB1
SUZ12
POU5F1
NANOG
TET1
Dr1
Mark
Prom CpG- .hg
Prom CpG+.hg
Promoters .hg
All regions .hg
Prom CpG- .mm
Prom CpG+.mm
Promoters .mm
All regions .mm
fold
3.0
2.5
2.0
1.5
1.0
overlap
0.2
0.4
0.6
0.8
1.0
B
hg
mm
H3K27ac
H3K4me1
H3K4me2
H3K4me3
H3K79me2
H3K79me3
H4ac
H4K5ac
H2BK20ac
H3K14ac
H3K36ac
H4K12ac
H4K8ac
H3K122ac
H3K56ac
H3K9ac
H3K9K14ac
H2A,Z
H3
H3,3
H3K18ac
H4K16ac
macroH2A1
macroH2A1,2
H2A,XS139ph
H2AK119Ub
H3K27me3
H3K36me3
H3K9me2
H3K9me3
Prom CpG-
Prom CpG+
Promoters
All regions
Prom CpG-
Prom CpG+
Promoters
All regions
fold
3.0
2.5
2.0
1.5
1.0
overlap
0.2
0.4
0.6
0.8
1.0
active transcription
active promoters
transcription activation
activation&repression
repressed->active
repression
enhancer
DNA damage
polycomb repression
bivalent chromatin
C
human Prom CpG+
human Prom CpG-
HM
TF
HM
TF

**Supplementary Figure 2.** Patterns of transcription factors and histone marks enriched around all Deep-Z predicted Z-flipons in human and mouse genome. **A.** Enrichment of transcription factors around Z-flipons for the entire genome, promoters, CpG-promoters and non-CpG promoters. **B.** Enrichment of histone marks around Z-flipons. **C.** Share of enriched histone marks and transcription factors around Z-flipons for CpG- and non-CpG promoters.

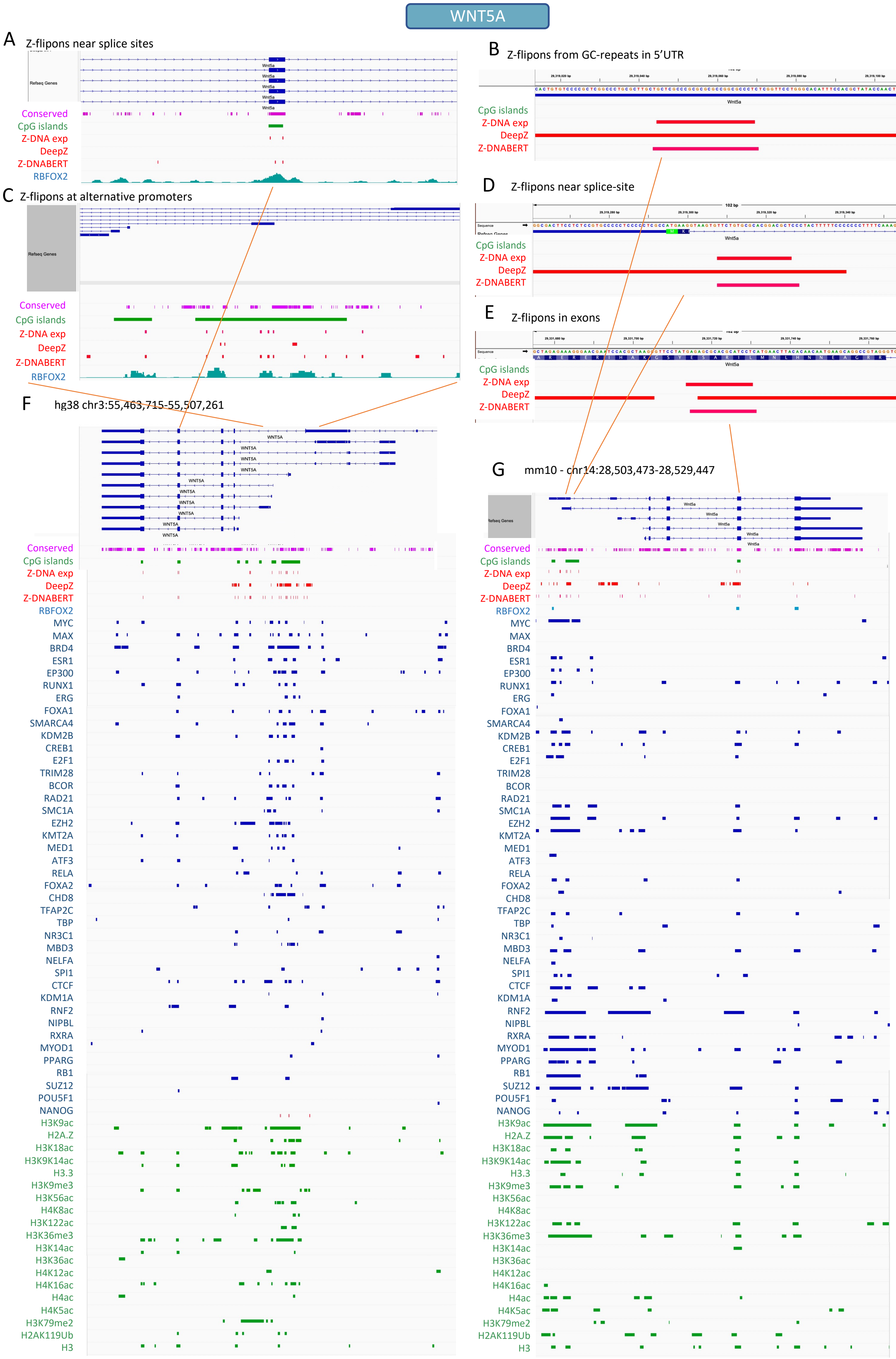
WNT5A
A Z-flipons near splice sites
Refseq Genes
Wnt5a
Conserved
CpG islands
Z-DNA exp
DeepZ
Z-DNABERT
RBFOX2
B Z-flipons from GC-repeats in 5'UTR
Wnt5a
CpG islands
Z-DNA exp
DeepZ
Z-DNABERT
C Z-flipons at alternative promoters
Refseq Genes
Conserved
CpG islands
Z-DNA exp
DeepZ
Z-DNABERT
RBFOX2
D Z-flipons near splice-site
102 bp
Sequence
Wnt5a
CpG islands
Z-DNA exp
DeepZ
Z-DNABERT
E Z-flipons in exons
Sequence
Wnt5a
CpG islands
Z-DNA exp
DeepZ
Z-DNABERT
F hg38 chr3:55,463,715-55,507,261
WNT5A
G mm10 - chr14:28,503,473-28,529,447
Refseq Genes
Wnt5a
Conserved
CpG islands
Z-DNA exp
DeepZ
Z-DNABERT
RBFOX2
MYC
MAX
BRD4
ESR1
EP300
RUNX1
ERG
FOXA1
SMARCA4
KDM2B
CREB1
E2F1
TRIM28
BCOR
RAD21
SMC1A
EZH2
KMT2A
MED1
ATF3
RELA
FOXA2
CHD8
TFAP2C
TBP
NR3C1
MBD3
NELFA
SPI1
CTCF
KDM1A
RNF2
NIPBL
RXRA
MYOD1
PPARG
RB1
SUZ12
POU5F1
NANOG
H3K9ac
H2A.Z
H3K18ac
H3K9K14ac
H3.3
H3K9me3
H3K56ac
H4K8ac
H3K122ac
H3K36me3
H3K14ac
H3K36ac
H4K12ac
H4K16ac
H4ac
H4K5ac
H3K79me2
H2AK119Ub
H3

**Supplementary Figure 3.** Z-flipons in WNT5A gene from Presynapse assembly pathway that was enriched in common cluster of conserved human and mouse Z-flipons (see Figure 3 in the main text). A-E. Regions of Z-flipons detected by three methods – DeepZ, KEx, Z-DNABERT. **A.** Conserved Z-flipons near splice sites of the second exon in WNT5A in human. **B.** Z-flipons composed from GC-repeats in 5'UTR region in mouse. **C**. Z-flipons at alternative promoters of WNT5A in human. **D**. Z-flipons near splice-site after the first exon of Wnt5a in mouse. **E**. Z-flipon detected by three methods – DeepZ, KEx, Z-DNABERT in the exon of Wnt5a in mouse. **F-G**. Region of WNT5A gene with signals from omics features enriched in Z-flipons both in human (**F**) and mouse (**G**) genomes. Omics features are aggregated signals from all tissues as they were used in DeepZ model.

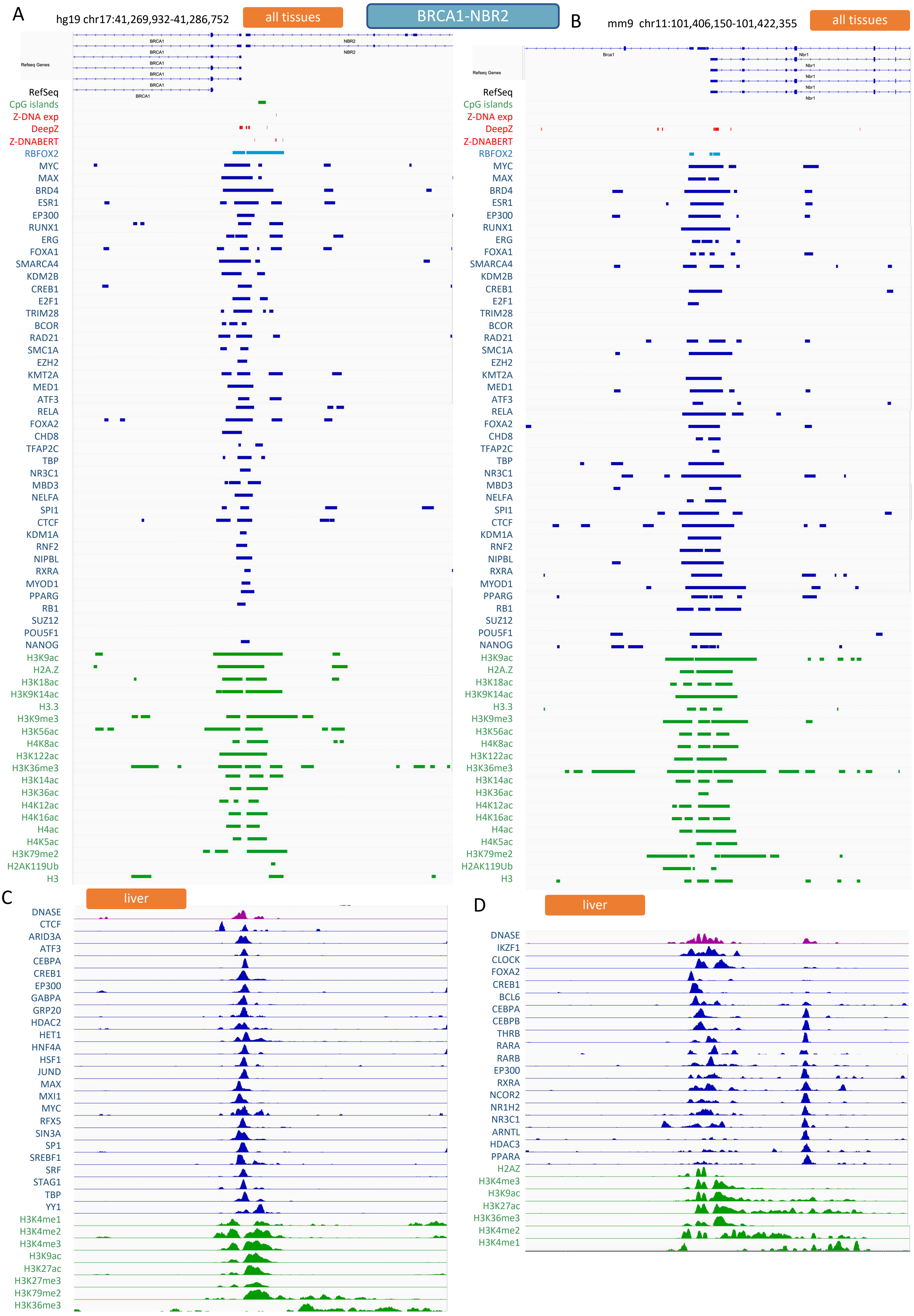
A
hg19 chr17:41,269,932-41,286,752
all tissues
BRCA1-NBR2
B
mm9 chr11:101,406,150-101,422,355
all tissues
Refseq Genes
BRCA1
NBR2
Brca1
Nbr1
RefSeq
CpG islands
Z-DNA exp
DeepZ
Z-DNABERT
RBFOX2
MYC
MAX
BRD4
ESR1
EP300
RUNX1
ERG
FOXA1
SMARCA4
KDM2B
CREB1
E2F1
TRIM28
BCOR
RAD21
SMC1A
EZH2
KMT2A
MED1
ATF3
RELA
FOXA2
CHD8
TFAP2C
TBP
NR3C1
MBD3
NELFA
SPI1
CTCF
KDM1A
RNF2
NIPBL
RXRA
MYOD1
PPARG
RB1
SUZ12
POU5F1
NANOG
H3K9ac
H2A.Z
H3K18ac
H3K9K14ac
H3.3
H3K9me3
H3K56ac
H4K8ac
H3K122ac
H3K36me3
H3K14ac
H3K36ac
H4K12ac
H4K16ac
H4ac
H4K5ac
H3K79me2
H2AK119Ub
H3
C
liver
DNASE
CTCF
ARID3A
ATF3
CEBPA
CREB1
EP300
GABPA
GRP20
HDAC2
HET1
HNF4A
HSF1
JUND
MAX
MXI1
MYC
RFX5
SIN3A
SP1
SREBF1
SRF
STAG1
TBP
YY1
H3K4me1
H3K4me2
H3K4me3
H3K9ac
H3K27ac
H3K27me3
H3K79me2
H3K36me3
D
liver
DNASE
IKZF1
CLOCK
FOXA2
CREB1
BCL6
CEBPA
CEBPB
THRB
RARA
RARB
EP300
RXRA
NCOR2
NR1H2
NR3C1
ARNTL
HDAC3
PPARA
H2AZ
H3K4me3
H3K9ac
H3K27ac
H3K36me3
H3K4me2
H3K4me1

**Supplementary Figure 4.** Z-flipons at bidirectional and alternative promoters of BRCA1-NBR2 in human and mouse genomes. **A-B.** Region around bidirectional promoter of BRCA1-NBR2 and signals from omics features enriched in Z-flipons both in human (**A**) and mouse (**B**) genomes. Omics features are aggregated signals from all tissues as they were used in DeepZ model. **C-D**. Selected omics features for human (**C**) and mouse (**D**) genome for liver tissue type.
