## Supplementary Table 2 for "Z-Flipons conserved between human and mouse are associated with increased transcription initiation rates"

**Supplementary Table 2.** Number of DeepZ predicted regions in human (HG) and mouse (MM) genomes depending on the quantile threshold. We chose the 3^rd^ quantile (marked in bold) according to PR curve that produced the best F-metric.

| **Quantile** | **Human predicted**  **(# regions)** | **Mouse predicted**  **(# regions)** | **Human**  **predicted**  **(length in bp)** | **Mouse**  **predicted**  **(length in bp)** | **Human**  **conserv**  **(length in bp)** | **Mouse**  **conserv**  **(length in bp)** |
| --- | --- | --- | --- | --- | --- | --- |

| 0.9 | 1 671 186 | 1 445 552 | 309 460 065  (10%) | 259 137 719  (10%) | 28 632 215  (9%) | 15 868 284  (6%) |
| --- | --- | --- | --- | --- | --- | --- |
| 0.99 | 264 017 | 153 892 | 31 011 539  (1%) | 26 659 234  (1%) | 44 018 09  (14%) | 1 777 033  (7%) |
| **0.999** | **30 083** | **17 569** | **3 071 150**  **(0,1%)** | **2 648 075**  **(0,1%)** | **602 020**  **(20%)** | **279 811**  **(11%)** |
| 0.9999 | 3 056 | 1 588 | 306 950  (0,01%) | 265 470  (0,01%) | 73 732  (24%) | 24 499  (9%) |
| 0.99999 | 238 | 126 | 30 700  (0,001%) | 26 609  (0,001%) | 7 397  (24%) | 777  (3%) |
| 0.999999 | 34 | 19 | 3081  (0,0001%) | 2684  (0,0001%) | 696  (23%) | 317  (12%) |
