## Supplementary Table 4 for "Z-Flipons conserved between human and mouse are associated with increased transcription initiation rates"

**Supplementary Table 4.** Top categories from the enrichment analyses of human and mouse ortholog genes with conserved Z-flipons in GO categories (sorted by FDR; see Full list of categories and genes in Supplementary Table 5).

| #category | term ID | term description | observed Nu gene | bgrnd Nugene | strength | FDR |
| --- | --- | --- | --- | --- | --- | --- |

| GO Process | GO:0019222 | Regulation of metabolic process | 546 | 6948 | 0,21 | 4,50E-36 |
| --- | --- | --- | --- | --- | --- | --- |
| GO Process | GO:0060255 | Regulation of macromolecule metabolic process | 518 | 6407 | 0,22 | 4,50E-36 |
| GO Process | GO:0080090 | Regulation of primary metabolic process | 490 | 6032 | 0,22 | 9,36E-34 |
| GO Process | GO:0010468 | Regulation of gene expression | 421 | 4813 | 0,25 | 1,11E-33 |
| GO Process | GO:0051171 | Regulation of nitrogen compound metabolic process | 478 | 5836 | 0,22 | 1,60E-33 |
| GO Process | GO:0031323 | Regulation of cellular metabolic process | 498 | 6239 | 0,21 | 6,23E-33 |
| GO Process | GO:0048522 | Positive regulation of cellular process | 453 | 5579 | 0,22 | 6,02E-30 |
| GO Process | GO:0009892 | Negative regulation of metabolic process | 307 | 3124 | 0,3 | 6,39E-30 |
| GO Process | GO:0019219 | Regulation of nucleobase-containing compound metabolic process | 360 | 3982 | 0,27 | 8,02E-30 |
| GO Process | GO:0010605 | Negative regulation of macromolecule metabolic process | 290 | 2875 | 0,31 | 1,07E-29 |
| GO Process | GO:0051252 | Regulation of RNA metabolic process | 340 | 3722 | 0,27 | 1,94E-28 |
| GO Process | GO:0009889 | Regulation of biosynthetic process | 369 | 4210 | 0,25 | 2,50E-28 |
| GO Process | GO:0010556 | Regulation of macromolecule biosynthetic process | 355 | 3976 | 0,26 | 2,50E-28 |
| GO Process | GO:0048518 | Positive regulation of biological process | 475 | 6112 | 0,2 | 6,66E-28 |
| GO Process | GO:0009893 | Positive regulation of metabolic process | 348 | 3893 | 0,26 | 9,69E-28 |
| GO Process | GO:0031326 | Regulation of cellular biosynthetic process | 361 | 4125 | 0,25 | 1,87E-27 |
| GO Process | GO:0010629 | Negative regulation of gene expression | 225 | 2014 | 0,36 | 2,23E-27 |
| GO Process | GO:2000112 | Regulation of cellular macromolecule biosynthetic process | 343 | 3878 | 0,26 | 1,98E-26 |
| GO Process | GO:0010604 | Positive regulation of macromolecule metabolic process | 326 | 3600 | 0,27 | 2,02E-26 |
| GO Process | GO:0031325 | Positive regulation of cellular metabolic process | 314 | 3413 | 0,27 | 2,76E-26 |
| GO Process | GO:0050789 | Regulation of biological process | 727 | 11475 | 0,11 | 1,76E-25 |
| GO Process | GO:0031324 | Negative regulation of cellular metabolic process | 261 | 2630 | 0,31 | 2,99E-25 |
| GO Process | GO:0051173 | Positive regulation of nitrogen compound metabolic process | 300 | 3239 | 0,28 | 2,99E-25 |
| GO Process | GO:0048519 | Negative regulation of biological process | 426 | 5389 | 0,21 | 3,54E-25 |
| GO Process | GO:0045935 | Positive regulation of nucleobase-containing compound metabolic process | 212 | 1927 | 0,35 | 7,95E-25 |
| GO Component | GO:0005634 | Nucleus | 590 | 7390 | 0,21 | 6,67E-45 |
| GO Component | GO:0005654 | Nucleoplasm | 381 | 3973 | 0,29 | 4,49E-38 |
| GO Component | GO:0031981 | Nuclear lumen | 423 | 4733 | 0,26 | 6,90E-37 |
| GO Component | GO:0005622 | Intracellular | 862 | 14276 | 0,09 | 2,19E-35 |
| GO Component | GO:0043229 | Intracellular organelle | 796 | 12528 | 0,11 | 3,23E-35 |
| GO Component | GO:0043231 | Intracellular membrane-bounded organelle | 721 | 10761 | 0,14 | 7,73E-35 |
| GO Component | GO:0043227 | Membrane-bounded organelle | 784 | 12427 | 0,11 | 4,48E-32 |
| GO Component | GO:0070013 | Intracellular organelle lumen | 470 | 5857 | 0,22 | 1,89E-31 |
| GO Component | GO:0043226 | Organelle | 821 | 13515 | 0,09 | 7,80E-30 |
| GO Function | GO:0003676 | Nucleic acid binding | 355 | 3947 | 0,26 | 1,87E-28 |
| GO Function | GO:1901363 | Heterocyclic compound binding | 459 | 5831 | 0,21 | 2,03E-27 |
| GO Function | GO:0097159 | Organic cyclic compound binding | 462 | 5916 | 0,2 | 4,50E-27 |
| KEGG | hsa05200 | Pathways in cancer | 64 | 517 | 0,4 | 5,19E-08 |
| KEGG | hsa05166 | Human T-cell leukemia virus 1 infection | 37 | 211 | 0,55 | 5,66E-08 |
| KEGG | hsa05225 | Hepatocellular carcinoma | 32 | 160 | 0,61 | 5,66E-08 |
| KEGG | hsa05205 | Proteoglycans in cancer | 34 | 196 | 0,55 | 1,71E-07 |
| KEGG | hsa05210 | Colorectal cancer | 22 | 82 | 0,74 | 1,71E-07 |
| KEGG | hsa05220 | Chronic myeloid leukemia | 21 | 75 | 0,76 | 1,71E-07 |
| KEGG | hsa05206 | MicroRNAs in cancer | 30 | 160 | 0,58 | 2,15E-07 |
| KEGG | hsa05213 | Endometrial cancer | 18 | 57 | 0,81 | 3,01E-07 |
| KEGG | hsa05226 | Gastric cancer | 28 | 144 | 0,6 | 3,01E-07 |
| KEGG | hsa04520 | Adherens junction | 19 | 67 | 0,76 | 4,02E-07 |
| KEGG | hsa04218 | Cellular senescence | 28 | 150 | 0,58 | 4,84E-07 |
| COMPARTMENTS | GOCC:0005634 | Nucleus | 444 | 4636 | 0,29 | 6,55E-47 |
| COMPARTMENTS | GOCC:0005622 | Intracellular | 767 | 11202 | 0,15 | 1,17E-45 |
| COMPARTMENTS | GOCC:0043229 | Intracellular organelle | 661 | 9242 | 0,17 | 9,91E-38 |
| COMPARTMENTS | GOCC:0043226 | Organelle | 684 | 9848 | 0,15 | 5,71E-36 |
| COMPARTMENTS | GOCC:0043231 | Intracellular membrane-bounded organelle | 563 | 7476 | 0,19 | 1,08E-33 |
| COMPARTMENTS | GOCC:0043227 | Membrane-bounded organelle | 617 | 8685 | 0,16 | 1,01E-31 |
| COMPARTMENTS | GOCC:0005654 | Nucleoplasm | 152 | 1110 | 0,45 | 1,54E-25 |
| COMPARTMENTS | GOCC:0031981 | Nuclear lumen | 195 | 1791 | 0,35 | 3,25E-22 |
